# A second wave of Notch signaling diversifies the intestinal secretory lineage

**DOI:** 10.1101/2024.07.15.603542

**Authors:** Eleanor Zagoren, Nicolas Dias, Zachary D. Smith, Nadia A. Ameen, Kaelyn Sumigray

## Abstract

The small intestine is well known for the function of its nutrient-absorbing enterocytes; yet equally critical for the maintenance of homeostasis is a diverse set of secretory cells, all of which are presumed to differentiate from the same intestinal stem cell. Despite major roles in intestinal function and health, understanding how the full spectrum of secretory cell types arises remains a longstanding challenge, largely due to their comparative rarity. Here, we investigate the fate specification of a rare and distinct population of small intestinal epithelial cells found in rats and humans but not mice: <u>C</u>FTR <u>Hi</u>gh <u>E</u>xpressers (CHEs). We use pseudotime trajectory analysis of single-cell RNA-seq data from rat intestinal jejunum to provide evidence that CHEs are specified along the secretory lineage and appear to employ a second wave of Notch-based signal transduction to distinguish these cells from other secretory cell types. We further validate the general order of transcription factors that direct these cells from unspecified progenitors within the crypt and experimentally demonstrate that Notch signaling is necessary to induce CHE fate both *in vivo* and *in vitro*. Our results suggest a model in which Notch is reactivated along the secretory lineage to specify the CHE population: a rare secretory cell type with putative functions in localized coordination of luminal pH and direct relevance to cystic fibrosis pathophysiology.

## Introduction

The physiological function of an epithelial tissue directly depends on the frequency and spatial distribution of its differentiated cell types, properties that are meticulously maintained by dedicated stem cell populations. In the mammalian small intestine, the epithelium is completely renewed every 3-7 days throughout adulthood (Barker et al., 2007), experiencing substantial cellular turnover without compromising its tissue morphology or function. Specifically, stem cells localized to crypt compartments continuously give rise to terminally differentiated cells that occupy the villus.

The primary role of the small intestine is to absorb nutrients from digested food. The function of secretion, however, is equally important for homeostasis, as it allows the intestinal epithelium to sense its highly dynamic luminal environment and release various ions, fluids and molecules to precisely modify its surroundings. These key functions are facilitated by specialized cell types. Intestinal epithelial differentiation is tightly regulated along two distinct cell fate lineages: absorptive and secretory. While absorptive cells are exclusively enterocytes, secretory cell types are varied and include anti-microbial Paneth cells, mucus-secreting Goblet cells, hormone-secreting Enteroendocrine cells (EECs), and chemosensory Tuft cells (Gehart and Clevers, 2019).

The diverse secretory lineage is thought to arise from a single secretory progenitor (Yan et al., 2012; Yang et al., 2001). The fundamental determinant of whether a cell will become an absorptive or secretory progenitor is Notch signaling via lateral inhibition, with absorptive progenitors “Notch-on” and secretory progenitors “Notch-off” (Fre et al., 2005; Yang *et al*., 2001). In the absorptive progenitor, Notch receptors regulate cell fate by promoting expression of the transcription factor Hes1, which initiates an extensive transcriptional program upon Notch activation. Conversely, in the secretory progenitor, Notch ligands induce a “Notch off” state, promoting the expression of the transcription factor Atoh1 to establish a secretory identity (Lo et al., 2017). Atoh1 has been described to be the master regulator of the secretory lineage, and it and Hes1 demonstrate reciprocal repression to reinforce cell fate decisions along their respective lineages (Yang *et al*., 2001). Loss of Atoh1 has been described to result in decreased populations of Goblet cells, Paneth cells, and EECs (Yang *et al*., 2001), although the latter group is exceptionally diverse in terms of cell subtypes (Haber et al., 2017) and has also been described to be specified in an Atoh1-independent manner (Gracz et al., 2018). Finally, the rare chemosensory population of Tuft cells remains one of the least well understood intestinal secretory cell types in terms of lineage dynamics, although they are thought to be specified independently of Atoh1 (Bjerknes et al., 2012; Gracz *et al*., 2018) and other major secretory transcription factors.

A less appreciated intestinal cell type with a putative secretory role and yet to be integrated into existing models of intestinal cell fate specification is the Best4+/<u>C</u>FTR <u>H</u>igh <u>E</u>xpresser (CHE). While present in humans (Burclaff et al., 2022; Busslinger et al., 2021; Strong et al., 1994), rats (Ameen et al., 1995), fish (Hiroi et al., 2005) and other vertebrates, an analogous cell type has not been found within the mouse intestine. As much of our knowledge of cell fate specification in the intestine has been informed by lineage tracing in genetic mouse models, we have a limited understanding of the mechanisms guiding CHE specification. Furthermore, the lack of tools to perturb CHE fate specification has precluded our ability to enrich for these cells and test their physiological contribution within the intestine. Although high expression of the chloride and bicarbonate ion channel CFTR (cystic fibrosis transmembrane conductance regulator)(Ameen *et al*., 1995) in CHEs hints at the capacity for intense secretory activity and a role in pH regulation, the function of CHEs and the mechanisms by which they are specified remain unknown.

In this study, we aim to explore CHE fate specification and integrate CHEs into the broader hierarchy of secretory cells. Specifically, we use pseudotime analysis of single cell RNA-seq data from rat intestinal jejunum to provide evidence that CHEs are specified along the secretory lineage. We further validate a set of distinguishing transcription factors and additional proteins to describe the biology of these cells from their initial induction within the crypt to their maturation within the villus. Unexpectedly, we identify an unusual reactivation of specific Notch receptors and effector proteins that strongly suggest the repurposing of this pathway to support CHE fate from a putative CHE-Tuft progenitor state. Together, our work reveals that active Notch signaling is necessary to induce CHE fate. This paradoxical reactivation of Notch signaling within the secretory lineage broadens our appreciation for the dynamic regulatory processes that diversify complex tissues.

## Results

### Rat CFTR High Expressers (CHEs) are derived from the secretory lineage

CHEs have been previously characterized as regionally restricted, cystic fibrosis transmembrane conductance regulator (CFTR)-positive cells within rat and human duodenum and proximal jejunum, but they are strikingly absent from the mouse intestine (Ameen *et al*., 1995; Busslinger *et al*., 2021; Strong *et al*., 1994). We validated the presence of the CHE population in the rat *in vivo*, where CFTR was highly enriched in CHEs in upper crypts and differentiated villi (Fig. 1A). In contrast, CFTR expression in mouse intestine was highest within crypts and apparent at low levels in the villus brush border, but not within discrete cell populations (Fig. 1B). Although CHEs have been hypothesized to regulate fluid secretion and/or luminal pH, their functional contributions to intestinal physiology have not yet been tested, in part because we lack understanding of how they are specified.

**Figure 1.**
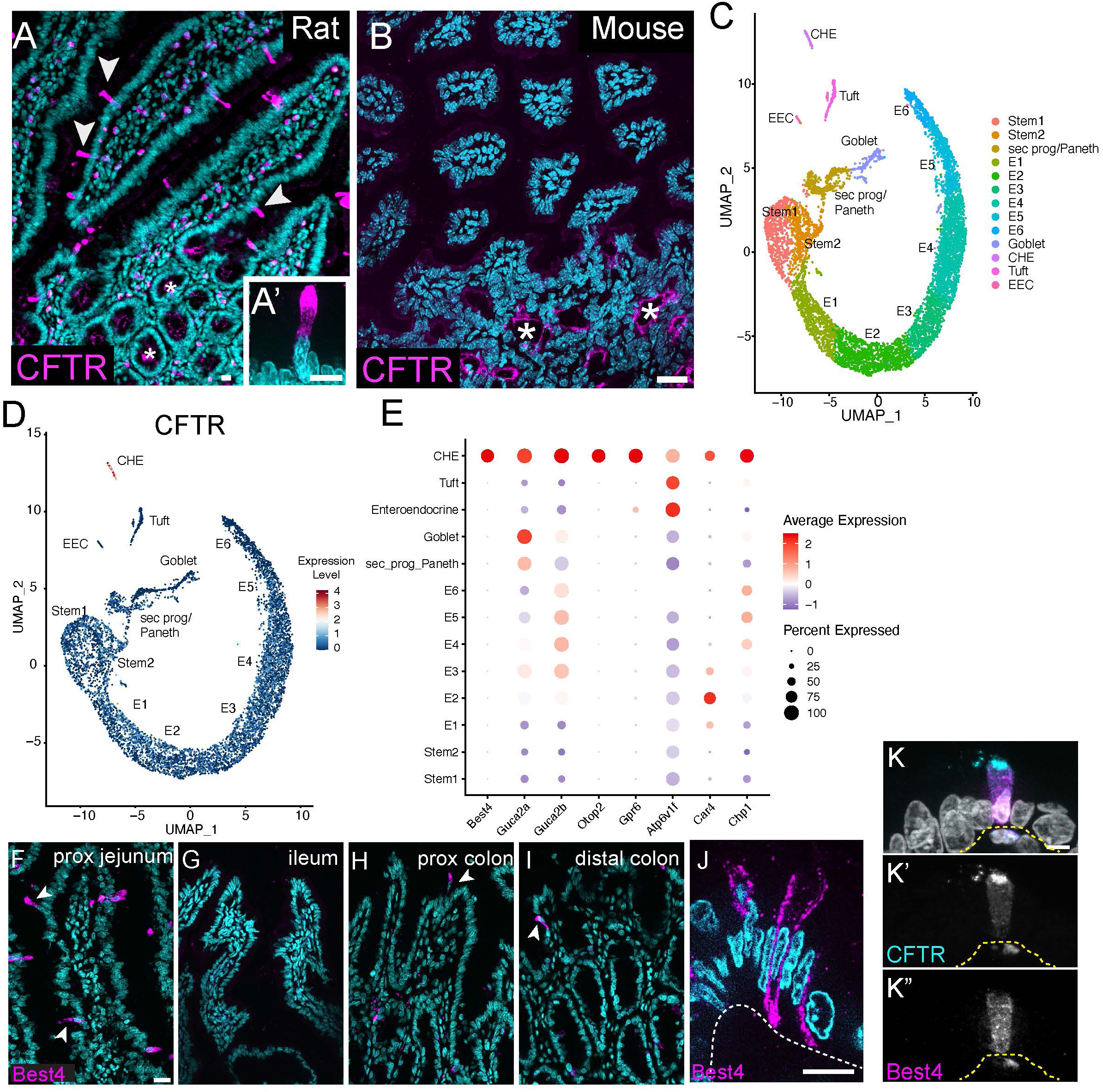
CHEs are a transcriptionally distinct intestinal epithelial cell type in rats but not mice. A) Rat intestinal jejunum stained for CFTR (magenta). Arrowheads denote CHEs in differentiated villi. * indicates CFTR enriched in crypts. DAPI in cyan. A’) CHE at higher magnification (63x). B) Mouse intestinal jejunum lacks CHEs when stained for CFTR (magenta). * indicates CFTR enriched in crypts. C) UMAP of rat intestinal jejunum epithelial cells (see Fig.S1 A-H for clustering annotation). E1-6 denote enterocytes at increasing stages of maturation. D) UMAP color coded for CFTR expression across intestinal epithelial cell types. E) Dotplot from scRNA-seq data demonstrating enrichment for proton-sensitive genes associated with pH regulation and ion and fluid homeostasis across epithelial cell types. F-I) Bestrophin 4+ cells (Best4, magenta) in rat proximal jejunum (F), rat ileum (G), rat proximal colon (H) and rat distal colon (I). DAPI in cyan. J) High magnification (63x) image of a single Z-slice of a Best4+ cell. Best4 in magenta, DAPI in cyan. Note basolateral Best4 localization. K) Co-staining of rat proximal jejunum for CFTR (cyan, K’) and Best4 (magenta, K’’). DAPI in gray. All scale bars, 10 μm

To characterize CHE gene expression more deeply, we performed unbiased transcriptomic analysis with droplet-based 3’ single-cell RNA sequencing (scRNA-seq) on 7,362 jejunal epithelial cells from two Sprague-Dawley rats. Clusters were annotated using known marker gene expression as a means of distinguishing distinct states (Fig. 1C and Fig. S1A-H). We first defined CHEs by their high *CFTR* expression (Fig. 1D) and confirmed that they did not express markers of other differentiated cell types or enterocyte-associated genes, as previously described by immunofluorescence (Ameen *et al*., 1995; Ameen et al., 2000; Jakab et al., 2013) (Fig. S1I). Instead, CHEs notably expressed multiple proton-sensitive genes implicated in pH regulation and ion and fluid homeostasis, including the bicarbonate transporter *Best4*, peptide hormones *Guca2a* (encodes guanylin) and *Guca2b* (encodes uroguanylin) and the ion channel *Otop2* (Otopetrin 2) (Fig. 1E).

These data are consistent with rat CHEs being analogous to the Best4+ “BCHE” population recently characterized in the human proximal small intestine (Burclaff *et al*., 2022; Busslinger *et al*., 2021). We confirmed that Best4+ cells were present in the rat proximal small intestine and colon, but not in the ileum (Fig. 1F-I), consistent with CHE localization. High magnification images of single slices of Best4+ cells in proximal jejunum demonstrate its basolateral membrane localization (Fig. 1J). Co-labeling of Best4 and CFTR in rat proximal small intestine confirmed that Best4+ cells in the small intestine and CHEs are indeed the same cell type (Fig. 1K), with CFTR enriched in the apical and subapical domains (Fig. 1K’) and Best4 enriched in the basolateral domains (Fig. 1K”). Taken together, CHEs broadly resemble a highly secretory cell type, including the specific trafficking of directional ion channels to modulate the intestinal lumen.

### CHEs have a distinct differentiation trajectory among secretory cells

Rat CHEs are functionally characterized as secretory cells (Jakab *et al*., 2013), and previous studies in human intestine suggest that small intestinal CHEs arise from secretory progenitors (Burclaff *et al*., 2022). In our data set, we observed a clear disparity between a densely sampled, continuous enterocyte trajectory and multiple disconnected clusters of differentiated secretory cells. Paneth and Goblet cells were obviously connected through a proliferative secretory progenitor. However, Tuft cells, EECs, and CHEs showed no obvious connectivity to each other nor to a distinct progenitor or stem population (Fig. 1C and Fig. S1J), making it unclear if CHEs derive from the secretory lineage or the absorptive lineage. Based on the human data (Burclaff *et al*., 2022), we hypothesized that CHEs arise from the secretory lineage. To test this, we generated diffusion maps of rat intestinal epithelial cells as a measure of pseudotime (Fig. 2A). Diffusion component 1 (DC1) largely captures differentiation of the absorptive lineage, while Diffusion component 2 (DC2) distinctly separates major secretory cell types, including CHEs.

**Figure 2.**
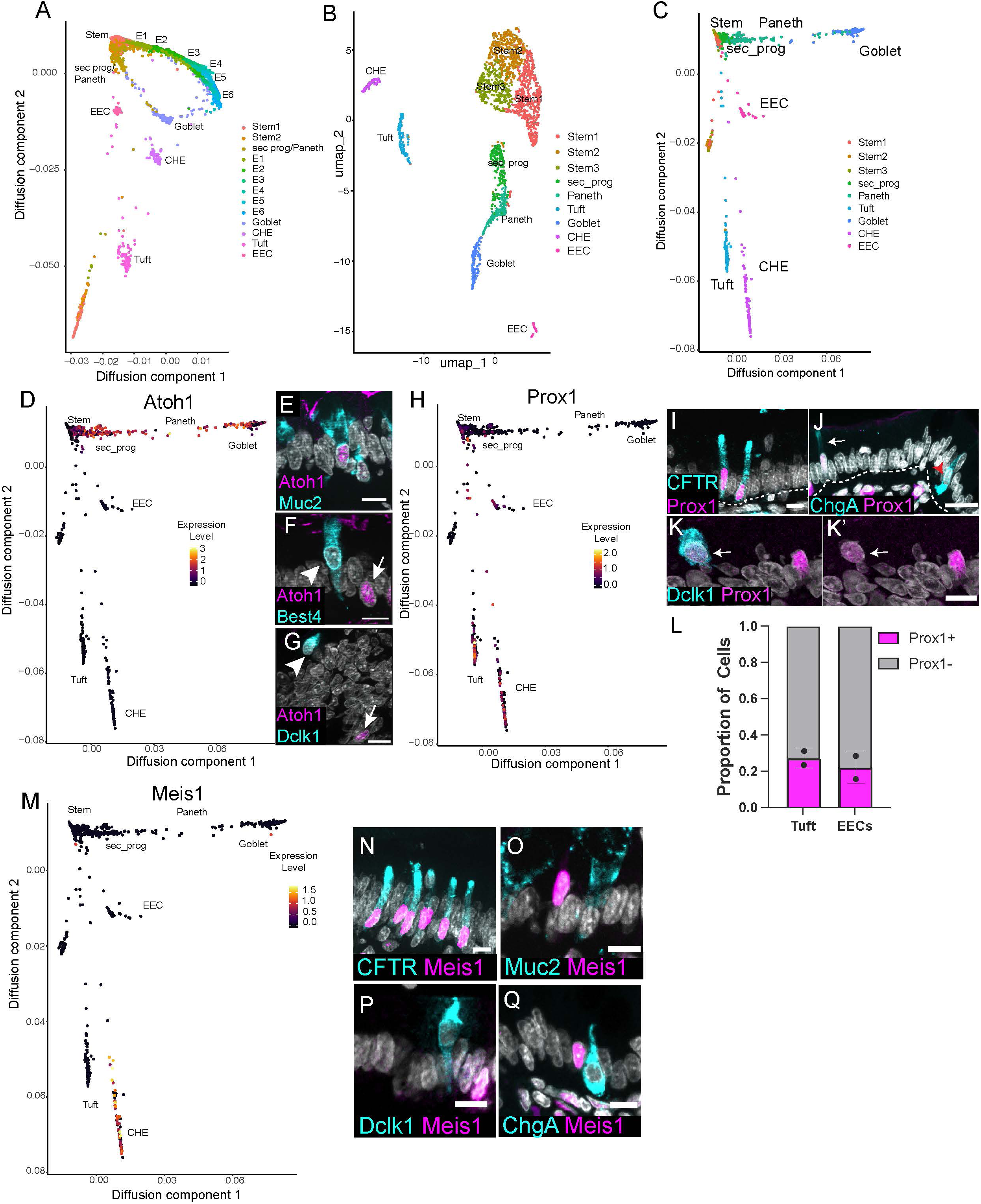
CHEs derive from a distinct branch of the secretory lineage shared with tuft cells and EECs and are enriched for the transcription factors Meis1 and Prox1. A) Diffusion maps of rat intestinal epithelial cells, showing their differentiation as a measure of pseudotime. E1-E6 denote enterocytes of increasing maturation. B) Reclustered UMAP of cells arising from stem and secretory progenitor clusters. C) Diffusion maps of stem cells and secretory progenitors as a readout of pseudotime. Note the clear bifurcation between Goblet/Paneth and the EEC/Tuft/CHE lineages. D) Diffusion map color coded for Atoh1 expression. E) Atoh1 (magenta) and Goblet cell staining (Muc2; cyan). F) Atoh1 (magenta) and Best4 (cyan; CHE marker) costaining. G) Atoh1 (magenta) and Dclk1 (cyan; Tuft cell marker) costaining. H) Diffusion map of stem and secretory cells color coded for Prox1 expression. I) CFTR (cyan; CHE marker) and Prox1 (magenta) co-staining. J) ChgA (cyan; EEC marker) and Prox1 (magenta) costaining. Arrow marks Prox1+/ChgA+ cell, red arrowhead marks Prox1-/ChgA+ cell. K-K’) Dclk1 (cyan, Tuft cell marker) and Prox1 costaining (in magenta, K’). L) Proportion of Tuft cells and EECs in rat proximal jejunum that are Prox1+. n=2 animals. Error bars, SD. M) Diffusion map of stem and secretory cells color-coded for Meis1 expression. N) Meis1 (magenta) and CFTR (cyan, CHE marker) costaining. DAPI in white. O-Q) Meis1 (magenta) staining with O) Muc2 (cyan, Goblet cells), P) Dclk1 (cyan, Tuft cells) or Q) ChgA (cyan, EECs). All scale bars, 10 μm.

To more sensitively examine the transcriptional relationships between distinct secretory cell types, we first subset our stem cells and secretory populations and reclustered our data (Fig. 2B, Fig. S2A-G). With the overpowering enterocyte populations removed, we were able to segregate a small secretory progenitor population, detectable by Dll1 expression (van Es et al., 2012) and cell cycle stage (Fig. S2H-I). Unlike the differentiated cells of the intestine, which were post-mitotic in G0, the secretory progenitor cluster contained cycling cells in both S and G2/M (Fig. S2I). *In vivo*, secretory progenitors have been described as mostly restricted to the crypt region above the uppermost Paneth cell and are marked by Dll1 expression (Sangiorgi and Capecchi, 2008; van Es *et al*., 2012; Yan *et al*., 2012). We also observed this population of secretory progenitors in the rat, which expressed high levels of Dll1 on the cell surface (Fig. S2J). Furthermore, these Dll1+ secretory progenitors were actively cycling, as evidenced by Ki67 expression (Fig. S2K).

We then performed pseudotime analysis of the stem cells and secretory cells by generating diffusion maps to identify clear sub-trajectories in their respective transcriptional behaviors. Interestingly, our diffusion map demonstrated a clear bifurcation between the Goblet/Paneth and the CHE/Tuft/EEC lineages (Fig. 2C), suggesting either the presence of multiple discrete progenitor cell types or a stepwise partitioning of a multipotent progenitor into more specialized states. To identify genes contributing to this bifurcation in downstream secretory cell types, we compared gene expression between the two groups, focusing particularly on transcription factors. As expected, the most highly enriched transcription factor along the Goblet/Paneth lineages was Atoh1 (Fig. 2D), a factor with a well-known role in consolidating a “Notch-off” state to support secretory cell differentiation (Yang *et al*., 2001). Immunofluorescence confirmed that Atoh1 protein was expressed in Goblet cells (Fig. 2E) but was absent from CHEs and Tuft cells (Fig. 2F-G). In contrast, the transcription factor Prox1 was the most highly enriched gene shared across the CHE/Tuft/EEC branch (Fig. 2H), suggesting two distinct secretory axes. Overall, our results reveal an intriguing bifurcation in transcriptional regulatory networks distinguishing CHE/Tuft/EECs from canonically Atoh1-positive Goblet/Paneth cells.

The restricted expression of Prox1 to CHEs, Tuft cells and EECs was particularly interesting, as the fly homolog of Prox1, Prospero, plays an important role in EEC fate in *Drosophila* adult midgut (Zeng and Hou, 2015). Prox1 has also been previously shown to mark some Tuft cells and EECs in mouse (Yan et al., 2017). Immunofluorescence of rat proximal jejunum further confirmed the expression of Prox1 in CHEs (Fig. 2I) and some EECs and Tuft cells (Fig. 2J-L). Notably, Prox1+ EECs had lower levels of the EEC marker chromogranin A (ChgA) compared to Prox1-/ChgA+ EECs (Fig. 2J). We did not detect a difference in Dclk1 expression in Tuft cells based on their expression of Prox1; however, all Tuft cells had lower Prox1 expression compared to CHEs (Fig. 2K-K’, arrow). The shared expression of Prox1 within a distinct sub-branch of the secretory lineage prompted us to search for additional factors that might support cell fates downstream of Prox1 induction. To understand how CHE fate is uniquely specified, we performed differential gene expression analysis of the CHE cluster compared to other cell states and manually curated a list of candidate transcription factors as likely drivers of cellular identity. Notably, *Meis1* was the sole transcription factor candidate exclusive to CHEs and not found in any other intestinal epithelial cell type (Fig. 2M, Fig. S2L). Immunofluorescence confirmed that Meis1 is specifically expressed in CHEs, marked by CFTR (Fig. 2N), and not in other secretory cell types, including Goblet cells (Fig. 2O), Tuft cells (Fig. 2P) or EECs (Fig. 2Q). We also examined Best4+ colon cells. While they have some overlapping gene expression with small intestinal CHEs (namely Best4 and Otop2), they do not express CFTR like small intestinal CHE/BCHEs (Burclaff *et al*., 2022). Surprisingly, while Best4+ colon cells were Prox1+ (Fig. S2M), Meis1 protein was undetectable (Fig. S2N), even though previous transcriptomic analysis had identified it as an enriched gene (Parikh et al., 2019; Busslinger *et al*., 2021; Burclaff et al., 2022). These results identify Meis1 as a novel CHE marker and suggest that it may play an important role in CHE differentiation and functional identity.

### CHEs express markers of active Notch signaling

Atoh1 is expressed in canonical secretory cells, where it reinforces the inhibition of Notch signaling and its associated transcriptional program (Lo *et al*., 2017; Yang *et al*., 2001). In contrast, Atoh1 expression is repressed in the absorptive lineage by the downstream Notch target Hes1 (Yang *et al*., 2001), making Notch signaling a well-established driver of the decision to become an absorptive or secretory progenitor (Gehart and Clevers, 2019; Yang *et al*., 2001). However, our data above demonstrate that the CHE/Tuft/EEC branch of the secretory lineage does not express Atoh1 (Fig. 2D-G), raising uncertainty about the role of Notch repression in maintaining cell fates for this branch of the secretory lineage. Therefore, we aimed to determine the Notch signaling status of cells within the CHE/Tuft/EEC branch.

Unexpectedly for a cell type arising from the secretory lineage, scRNA-seq analysis revealed that CHEs expressed factors associated with active Notch signaling, including the *Notch1* and *Notch2* receptor genes (Fig. 3A). More broadly, when we applied collective modules for “Notch-on” and “Notch-off” gene signatures as described in the Experimental Procedures, we observed that the “Notch on” signature was high within the CHE branch (Fig. 3B), while a clear “Notch off” signature was apparent in the Goblet/Paneth branch (Fig. 3C). Furthermore, at the transcriptomic level, our scRNA-seq analysis indicated that CHEs were the only cells in the rat intestine highly enriched for the *Notch2* receptor (Fig. 3A). We confirmed that CHEs specifically expressed the Notch2 receptor protein via immunofluorescence (Fig. 3D). Notch2 has been historically shown to specify cell types independently of the Notch1 receptor (Basil et al., 2022; Jeliazkova et al., 2013; Nakano et al., 2022), and we were intrigued by the possibility of a Notch2-specific role in CHE fate specification.

**Figure 3.**
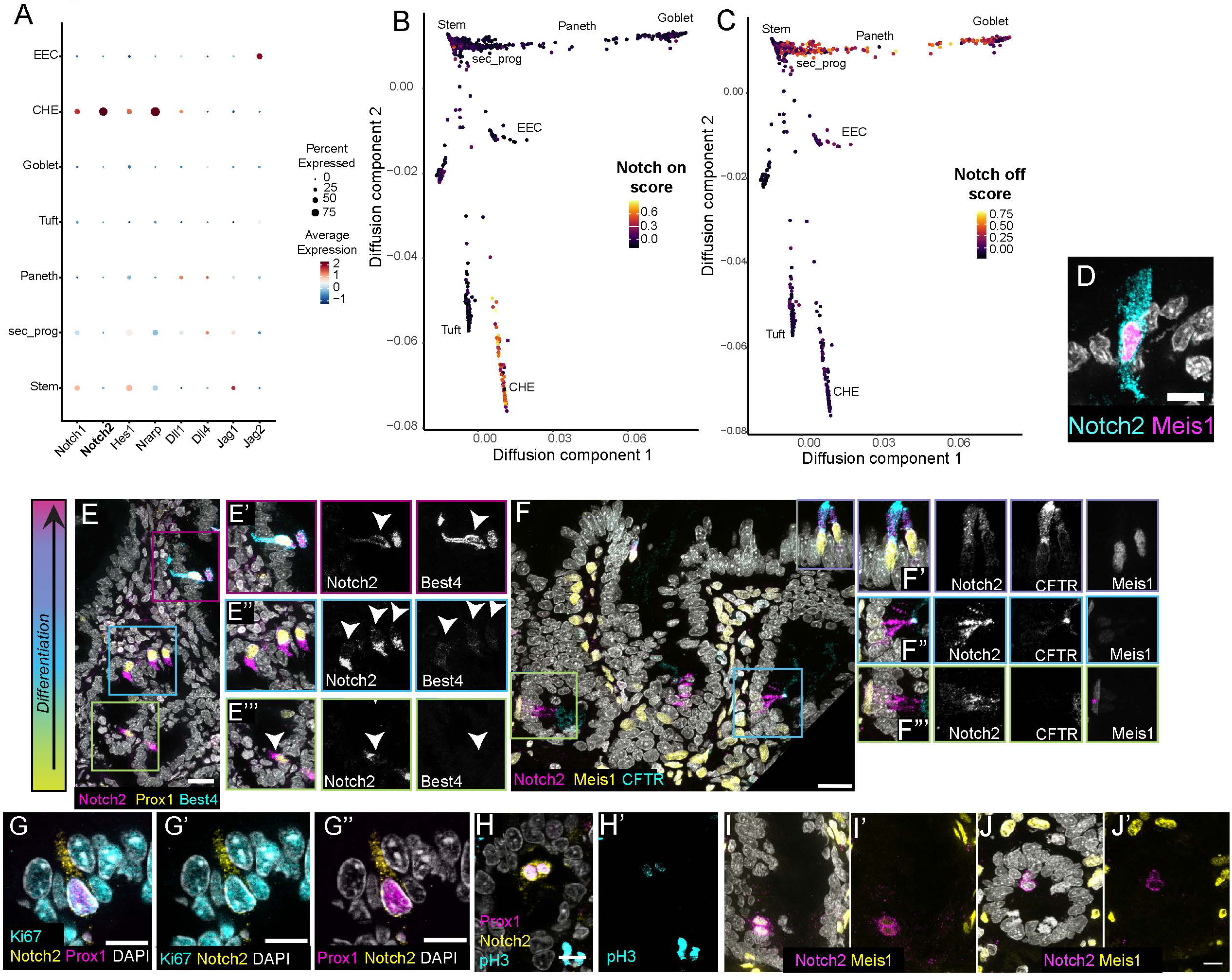
Notch2 is a candidate marker of CHE progenitors. A) Dotplot from scRNA-seq data showing relative expression of factors associated with Notch signaling across intestinal epithelial cell types. B-C) Module scores for Notch-on (B) and Notch-off (C) gene signatures. D) Meis1 (magenta, CHEs) and Notch2 receptor (cyan) costaining. DAPI in white. E) Rat proximal jejunum stained for Best4 (cyan), Prox1 (yellow), and Notch2 (magenta). E’) triple positive CHEs in the differentiated villus region, E’’-E’’’) Notch2+/Prox1+ cells in crypts that do not express Best4. Color of insert box denotes approximate localization along the crypt-villus axis (left). DAPI in white. F) Rat proximal jejunum stained for Notch2 (magenta) and Meis1 (yellow) with CFTR (cyan). Color of insert box denotes approximate localization along the crypt-villus axis (left; F’-F”’). In F”’, magenta asterisk denotes a stromal non-epithelial expressing Meis1. G-G”) Notch2+ /Prox1+ cell expressing Ki67. Notch2 in yellow, Prox1 in magenta, Ki67 in cyan. H-H’) Notch2+/Prox1+ crypt cells in active mitosis. Note symmetric cell division for both Notch2 (yellow) and Prox1 (magenta). H’) Low level of phospho-histone 3 (in cyan) as cell is exiting mitosis. I-I’) Notch2+/Meis1^Low^ cells in active mitosis. Notch2 (magenta), Meis1 (yellow), DAPI in white. J-J’) Notch2+ cell lacking Meis1 in active mitosis. Notch2 (magenta), Meis1 (yellow), DAPI in white. All scale bars, 10 μm.

### Notch2 is a candidate marker of CHE precursors

Intestinal epithelial cells progressively differentiate as they migrate upwards from the stem cell/progenitor zone in the crypt towards villi. In the absence of straightforward genetic lineage tracing tools in the rat, we took advantage of the position of cells along this progressively differentiating crypt-villus axis to infer the general spatial and temporal properties of CHE differentiation. We stained rat jejunum for CFTR along with our novel CHE markers to determine the temporal dynamics of CHE protein expression. We first examined Best4, Notch2, and Prox1 (Fig. 3E), and observed that as expected, CHEs in the differentiated villus region expressed high levels of Best4, Notch2 and Prox1 (Fig. 3E’). Similarly, labeling for CFTR, Meis1 and Notch2 (Fig. 3F) clearly identified triple-positive cells within the villus (Fig. 3F’).

Remarkably, we unexpectedly saw an additional population of Notch2+/Prox1+ cells in the proliferative crypt region that did not express Best4 (Fig. 3E”). These Notch2+ cells within the mid and upper crypts progressively expressed increasing levels of CFTR, Best4, and Meis1 before entering the villus compartment (Fig. 3E-F). These data are consistent with computational data from the human intestine which suggested that Best4 is a late mature CHE marker (Burclaff *et al*., 2022).

Interestingly, a Prox1+ progenitor has been previously described in the mouse with both reserve stem cell activity and rare but detectable clonogenic potential during homeostasis (Yan *et al*., 2017). This raises the intriguing hypothesis that the Notch2+/Prox1+/Best4-/CFTR-cells we observed in the mid-crypt region may function as a candidate progenitor, although it is also possible that these cells are simply immature CHEs that have not reached full differentiation status. To examine whether Notch2+/Prox1+ crypt cells in the rat could also act as a progenitor population, we first aimed to determine whether these cells were proliferative. We co-stained Notch2+/Prox1+ crypt cells for Ki67 and found that a subset of Notch2+/Prox1+ cells was Ki67+ (Fig. 3G-G”), suggesting that this population is still in the cell cycle and has not terminally differentiated and entered G0. In contrast, Ki67+ cells were not observed in differentiated villar CHEs (data not shown), and our scRNA-seq data indicated that differentiated CHEs fully exist within the G0/G1 stage of the cell cycle (Fig. S2I). While rare, we also detected Notch2+/Prox1+ crypt cells in active mitosis (Fig. 3H-H’). In these cases, Notch2 and Prox1 were expressed in both daughter cells, suggesting that Notch2+/Prox1+ cells undergo symmetric divisions. Finally, by examining Meis1 levels in mitotic cells, we found that some Notch2+ cell pairs were both Meis1+ (Fig. 3I-I’), while others were Meis1-(Fig. 3J-J’). We did not observe any instances in which dividing cells were asymmetrically Meis1+. These data suggest that Notch2+ cells are proliferative and arise within the crypt, and that rather than simply being immature CHEs, they may maintain the potential to generate other cell types within the CHE/Tuft/EEC axis.

### CHE fate specification requires active Notch signaling

While the presence of the Notch2 receptor strongly suggests active Notch signaling in CHEs, receptor expression does not necessarily reflect downstream Notch signaling activity. Therefore, we tested whether Hes1, the dominant effector transcription factor for active Notch signaling in the intestine, was expressed in CHEs. Co-staining of Hes1 and Meis1 revealed that CHEs expressed high levels of nuclear Hes1 (Fig 4A, arrows), strongly implying that CHEs engage in active Notch signaling. We also confirmed nuclear Hes1 signal in proliferative Meis1-crypt cells as a positive control (Fig.4A’, arrowheads), as the requirement for Notch signaling in intestinal stem cell homeostasis and transit amplifying cells has been well-characterized (Pellegrinet et al., 2011). Interestingly, the fluorescence intensity of nuclear Hes1 in CHEs was significantly higher than that of stem/TA cells (Fig. 4B), supporting the idea that Notch signal transduction is an active part of CHE differentiation. Notably, the nuclear Hes1 signal in the stem/progenitor zone were also considerably more heterogeneous compared to the stable Hes1 levels observed in CHEs (Fig. 4C), perhaps due to the oscillatory dynamics of this TF, which have been previously reported in other systems (Hirata et al., 2002; Oates et al., 2012; Seymour et al., 2020).

**Figure 4.**
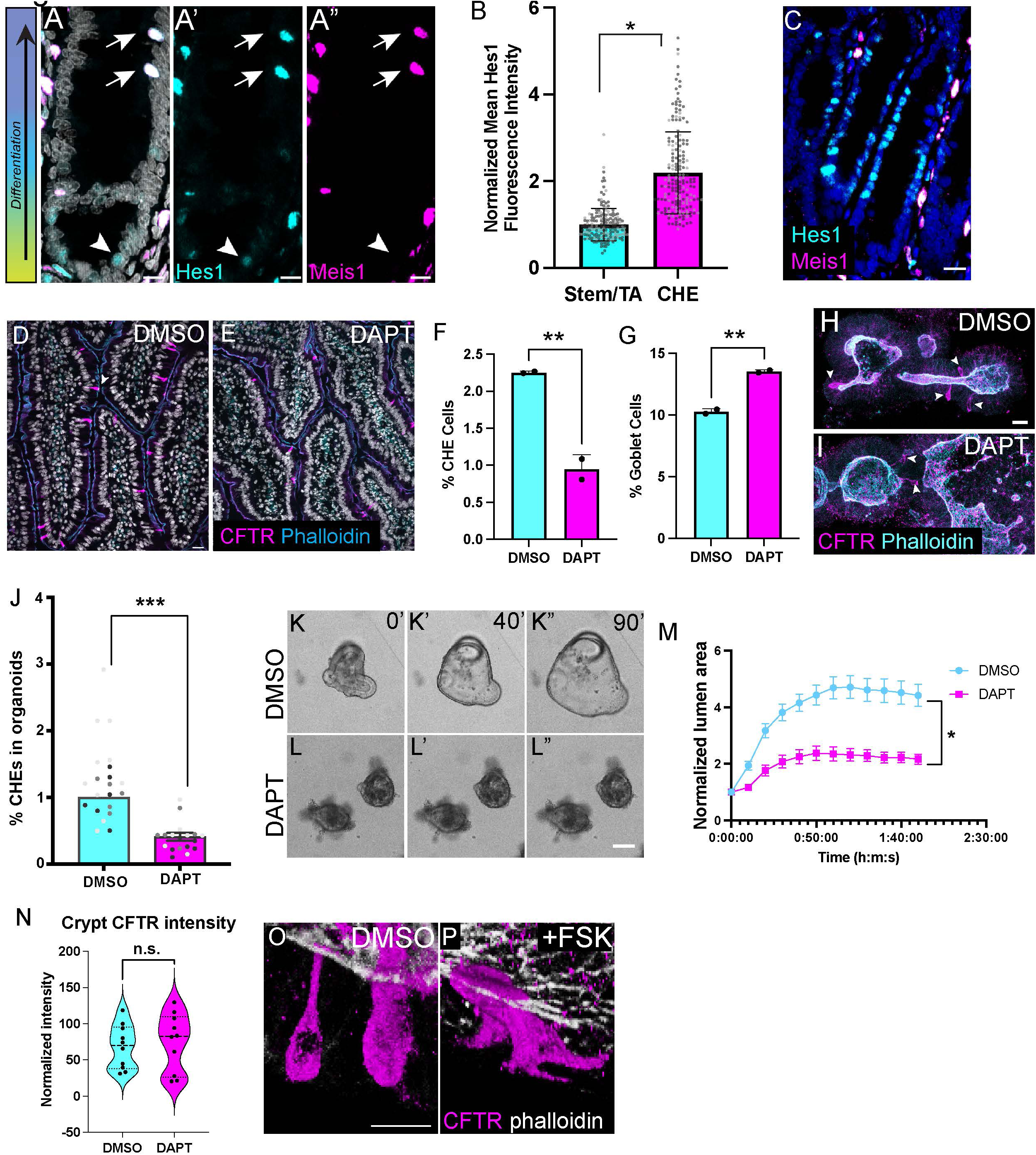
CHE specification requires active Notch signaling. A) Meis1 (magenta; CHE marker; arrowheads) and Hes1 (cyan) costaining. Stem/TA cells with positive Hes1 staining marked with an arrow. Scale bar, 10 μm. B) Fluorescence intensity of nuclear Hes1 in CHEs and stem/TA cells. n=2 rats, unpaired two-tailed t-test. *p*<0.01. Each point denotes one cell. Shade of gray indicates different technical replicates. Error bars indicate SD. C) A) Hes1 (cyan) fluorescence intensity levels in rat intestinal jejunum are heterogeneous within the crypt region. Scale bars, 20 μm. D-E) Immunofluorescence staining for CHEs (CFTR, magenta) and F-actin (cyan, phalloidin) for vehicle (D) and DAPT-treated rats (E). DAPI in white. Scale bar, 10 μm. F-G) Percentage of CHEs (F, *p*<0.01) and % Goblet cells (G, *p*<0.003) in DAPT-treated rats relative to DMSO controls. Two-tailed unpaired t-test. Error bars denote SD. H-I) Immunofluorescence staining for CHEs (CFTR, magenta) for control (H) and DAPT-treated (I) rat intestinal organoids. Arrowheads mark CHEs. Scale bar, 20 µm. J) Quantification of normalized CHE abundance in DMSO vs. DAPT-treated organoids. n=3 replicates, each point denotes one organoid. Two-tailed unpaired t-test. p<0.0006. Error bars denote SD. K-L) Forskolin swelling for control (K) and DAPT-treated (L) rat intestinal organoids. K-L) time=0, K’-L’) time=40 min, K’’-L’’) time =90 min. Scale bar, 100 μm. M) Normalized lumen area for DMSO (control) vs. DAPT-treated organoids over time following forskolin swelling. Unpaired t-test, area under the curve at 110 min for N=2 technical replicates. *p*<0.04. Error bars denote SD. N) Quantification of normalized crypt CFTR intensity for DMSO (cyan) and DAPT (magenta) treated organoids. *n.s.,* not significant. O-P) CHE morphology in organoids with DMSO treatment (O), and with forskolin (FSK) treatment (P). CFTR (magenta) marks CHEs, phalloidin (gray) strong marks the apical domain. Scale bar, 10 µm.

To test whether Notch signaling was necessary for CHE fate, we injected adult rats for five consecutive days with DAPT, a γ-secretase inhibitor that blocks the cleavage of the Notch intracellular domain, thereby inhibiting downstream Notch signaling. The experimental timeline accounts for the complete renewal of the intestinal epithelium every 3-7 days (Barker *et al*., 2007). DAPT treatment significantly decreased CHE number relative to the DMSO control, demonstrating that CHE fate specification requires active Notch signaling (Fig. 4D-F). Conversely, we found that Goblet cell number increased upon Notch inhibition (Fig. 4G), as expected from previous reports (Yang *et al*., 2001) and consistent with the relative positioning of CHE vs. Goblet cells according to our diffusion component mapping.

To more robustly test mechanisms driving fate specification of CHEs and ultimately, CHE function, we generated intestinal organoids from the rat jejunum (Zagoren et al., 2023). We first aimed to determine whether CHEs were specified in rat organoid models. Undifferentiated intestinal spheroids present two days after culturing did not contain CHEs (Fig. S3A-B). However, upon organoid budding and differentiation, CHEs were clearly defined (Fig. S3C-D). CHEs in organoids localized to the same developmental domains as found *in vivo*, including enrichment within the differentiated domains of the upper crypt and villus regions (Fig. S3E). Outside of CHEs, rat organoid enterocytes also expressed baseline CFTR in a gradient similar to that seen *in vivo*, with highest levels in the crypt and low levels in villar domains (Fig. S3F). CHEs in organoids also expressed Meis1, Prox1, Best4, and Notch2 as *in vivo* (Fig. S3G-J). We then verified whether Notch signaling regulates CHE fate *in vitro* as it does *in vivo*. We allowed organoids to differentiate, then treated them with 5 µM DAPT for three days. After DAPT treatment, we assessed the relative abundance of CHEs in DAPT vs. vehicle (DMSO) treatment. As found *in vivo*, DAPT treatment significantly inhibited CHE specification (Fig. 4H-J). These data suggest that active Notch signaling is required for CHE fate specification in organoids, as it is within intact tissue.

Given the relative ease with which we could chemically modulate CHE number within organoids, we next sought to evaluate a potential secretory function for CHEs and their contribution to fluid secretion. Forskolin, an adenylyl cyclase activator, is commonly used in organoid-based assays to test for the ability to swell and secrete fluid (Boj et al., 2017). Vehicle (DMSO) or DAPT-treated organoids were treated with forskolin to induce fluid secretion into the lumen, and the degree of organoid swelling was observed. Notably, decreased CHE specification in DAPT-treated organoids resulted in strong abrogation of organoid swelling (Fig. 4K-M), indicating a dramatic change in flux between the external medium and the luminal space and supporting a role for CHEs in sustained fluid secretion in intestinal organoids. However, Notch signaling is required for stem cell maintenance, and outside of CHEs, most CFTR is expressed within the crypt domain. To rule out the possibility that we were affecting both the CHE population and the crypt pool of CFTR, we measured the fluorescence intensity of CFTR within crypts of vehicle and DAPT-treated organoids. No significant difference was observed in normalized crypt CFTR intensity (Fig. 4N), suggesting that the secretory defects observed in DAPT-treated organoids are not attributable to CFTR loss in the crypt compartment but rather to reduced CHE numbers. Surprisingly, when we stained rat organoids for CFTR to examine CHEs after forskolin activation, we found that they had dramatically changed shape, forming long protrusions (Fig.4O-P). In fixed tissue *in vivo*, we could occasionally capture CHEs with long protrusions (data not shown), similar to what has been described for rare CFTR-enriched pulmonary ionocytes in the airway (Yuan et al., 2023) and suggesting that CHEs can sense and respond to their environment.

In summary, CHEs comprise a rare cell type in the proximal small intestine that likely arises from the secretory lineage along a distinct branch shared with Tuft cells and EECs. CHEs are specifically enriched for Meis1 and share expression of Prox1 with Tuft and EEC populations. Notch2 is a candidate marker of CHE progenitors, as we observe Notch2+/Prox1+ cells in crypts that lack enrichment for the CHE markers Best4 and CFTR. Finally, Notch signaling is necessary for CHE cell fate. These data support a model in which CHEs arise from the secretory lineage, yet paradoxically, Notch must be reactivated to specify this rare population (Fig. 5).

**Figure 5.**
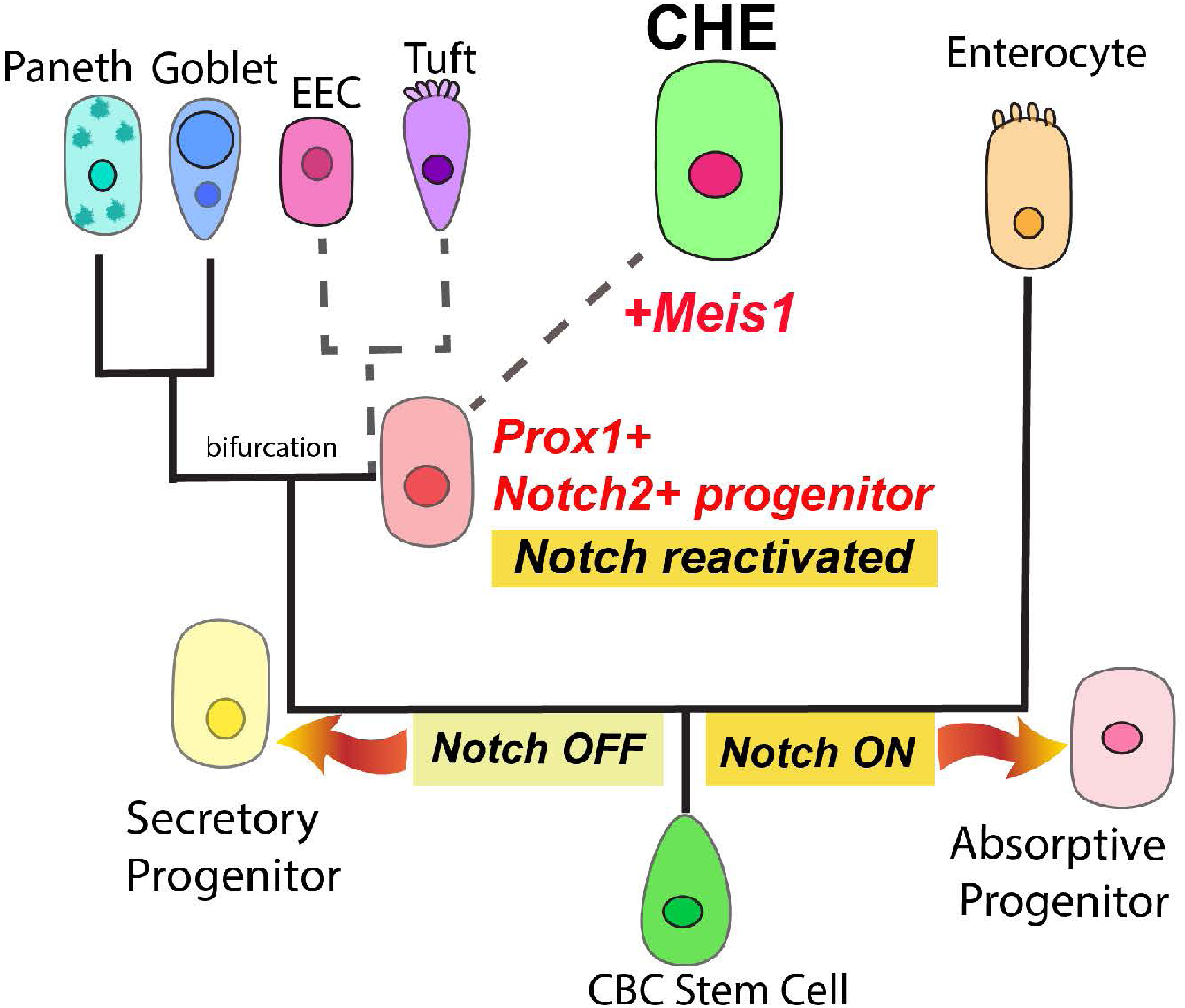
Working model for CHE cell fate specification. Working model of CHE cell differentiation in which: 1) CHEs arise from the secretory lineage, which is initially Notch off, 2) Notch is reactivated along the secretory lineage as a potentially multipotent Notch2+/Prox1+ progenitor and 3) Meis1 is upregulated to promote the expression of differentiated CHE markers, including CFTR and Best4.

## Discussion

In this study, we found that rare intestinal epithelial CHEs originate from the secretory lineage and depend on active Notch signaling for their fate specification. We used scRNA-seq from epithelial cells of rat jejunum and generated diffusion maps of stem and secretory cells to illustrate that CHEs likely arise from the secretory lineage. Furthermore, through pharmacological inhibition experiments, we demonstrated that Notch signaling is necessary for CHE cell fate specification both *in vivo* and *in vitro*.

These findings collectively suggest that Notch must be *reactivated* along the secretory lineage to drive CHE fate, likely via the induction of alternative receptors such as Notch2, and may have broader implications for the behavior and diversification of specialized secretory cells. In particular, this notion of Notch reactivation in CHEs challenges the prevailing model of the secretory lineage as “Notch off.” However, this model of a wholly “Notch off” secretory lineage has been informed by lineage tracing using mouse models (Yang *et al*., 2001), which do not generate a comparable CHE/BCHE population under homeostatic conditions. Moreover, it is challenging to appropriately evaluate the secretory lineage considering the comparative rarity and complexity of secretory cell types, even when conventional genetic lineage tracing techniques are available. Our evidence of Notch reactivation during CHE differentiation cautions against inferring the final downstream Notch signaling status in differentiated cell types from the initial Notch signaling status of their less determined progenitors. Alternate methods for transducing these signals can be temporally restricted to ensure an orderly transition in developmental potential. Ultimately, our results highlight the surprising modularity of the secretory lineage despite its restricted origin within the intestinal crypt.

We propose a working model of CHE cell differentiation in which: 1) CHEs arise from the secretory lineage, which is initially Notch off, 2) Notch is reactivated along the secretory lineage as a potentially multipotent Notch2+/Prox1+ progenitor and 3) Meis1 is upregulated to promote the expression of differentiated CHE markers, including CFTR and Best4. While we and others (Burclaff *et al*., 2022) have computationally described CHEs and BCHEs respectively as arising from the secretory lineage, our work further identifies a resident Notch2+/Prox1+ progenitor in crypts that can divide symmetrically, suggesting that Notch reactivation may precede terminal CHE differentiation.

Beyond their localization to the crypt, undifferentiated Notch2+/Prox1+ cells generally lack Meis1, the only CHE-specific transcription factor identified in our scRNA-seq, but can occasionally be found with low levels of nuclear Meis1 that suggests the onset of CHE differentiation. The full lineage generating capability of this Notch2+/Prox1+ progenitor *in vivo* remains to be tested and is currently limited by the lack of genetic tools available in rat and human models. Nevertheless, a previous mouse study characterized Prox1+ progenitors as having both reserve stem cell activity and rare clonogenic potential in homeostasis (Yan *et al*., 2017), features that our putative Notch2+/Prox1+ progenitor may share in the rat.

Finally, we have shown that the transcription factor Meis1 is specific to CHEs and is not present in any other intestinal epithelial cell type. As Meis1 is the only transcription factor candidate uniquely enriched in CHEs, it likely plays a key role in regulating small intestinal CHE fate, potentially as a terminal differentiation factor capable of upregulating the expression of ion channels required for luminal secretion such as CFTR and Best4. However, whether Meis1 itself is necessary for CHE fate has yet to be investigated, but our validation that these cells arise as part of intestinal organoid differentiation provides a new more tractable model to investigate this and other hypotheses. Collectively, these results highlight the value of exploring the secretory lineage beyond the characterization that has been done in mice. Furthermore, the increasing diversity of cell types that arise from the same secretory progenitor warrants detailed characterization of their gene regulatory networks, including the specific transcription factor binding profiles and stepwise order of molecular events required to appropriately partition and define their unique functions.

Intriguingly, our data reveal a high nuclear Hes1 intensity level in differentiated CHEs and considerably lower, heterogenous nuclear Hes1 levels in the stem/progenitor zone. Hes1 is notable for its oscillatory activity (Hirata *et al*., 2002), which regulates differentiation timing in a variety of systems (Maeda et al., 2023; Seymour *et al*., 2020). The behavior of Hes1 between stem/progenitor cells and CHEs may represent a transition from oscillatory to sustained Hes1 expression during CHE differentiation. Hes1 dynamics have been shown to impact proliferative ability, including recent work in neural stem cell culture where oscillating Hes1 levels in progenitors inhibit p21 to promote proliferation, and high sustained Hes1 levels upregulate p21 to promote quiescence (Maeda *et al*., 2023). It is enticing to hypothesize that CHEs may behave analogously to this model, with the transition from oscillatory to high sustained Hes1 signaling promoting their quiescence and terminal differentiation. Moreover, in the intestine, positional information provided by the Wnt gradient may also greatly inform oscillatory Notch dynamics to control the timing and positioning of distinct secretory cell types. Although modeled computationally (Kay et al., 2017), these dynamics have yet to be experimentally investigated *in vivo*.

The fact that CHEs are so highly enriched for Hes1 and require active Notch signaling for their fate specification is perhaps less surprising when considered alongside pulmonary ionocytes, a rare CFTR enriched cell type within the lung that also requires Notch signaling for proper specification (Plasschaert et al., 2018). Although these cell types appear analogous, there is no evidence of Foxi1 (Forkhead Box I1) expression within any intestinal cell type (data not shown), even though this transcription factor is necessary and sufficient for pulmonary ionocyte fate (Montoro et al., 2018; Plasschaert *et al*., 2018) and is expressed within the ionocytes of many tissues (Pou Casellas et al., 2023). Notably, Foxi1 has also been described as promoting ionocyte, Tuft cell, and neuroendocrine cell specification through a shared rare Foxi1+ progenitor in developing airway epithelium (Yuan *et al*., 2023). Although these airway cell types exhibit highly similar transcriptional relationships to those we have found for CHEs, Tufts and EECs within the intestinal secretory lineage, the total absence of Foxi1 within our small intestinal transcriptional atlas indicates that similar developmental relationships may hold despite an alternative transcriptional program. As a putative alternative, our results and the work of others (Yan *et al*., 2017) suggest that this unique secretory sub-branch may be driven more directly via Prox1, a possibility that warrants further characterization.

While ionocytes were initially described as a major source of CFTR activity and secretion in the lung (Plasschaert *et al*., 2018), subsequent work has clarified that the majority of CFTR expression and secretion is performed by other secretory cells, which vastly outnumber rare ionocytes (Okuda et al., 2021). Nevertheless, ionocytes have been hypothesized to be involved in the regional coordination of pH sensing and acute responses to hyperosmotic stress across a range of systems (Okuda *et al*., 2021; Yuan *et al*., 2023), and CHEs may similarly coordinate a regionalized pH regulation in response to lumen acidification. CHEs are highly enriched for apical CFTR, and thereby capable of bicarbonate efflux out into the luminal environment (Garcia et al., 2009). Furthermore, CHEs are regionalized to the distal duodenum and proximal jejunum, regions of the intestine where residual stomach acid must be effectively neutralized to promote optimal nutrient absorption (Jakab *et al*., 2013). This putative function would be particularly relevant for cystic fibrosis, where lumen acidification driven by the loss of CFTR-mediated bicarbonate secretion drives the gastrointestinal pathophysiology of cystic fibrosis and results in significant comorbidities (De Lisle and Borowitz, 2013; Garcia *et al*., 2009). Further studies will be needed to determine the specific functional contribution of CHEs to the overall physiology of the small intestine in the context of homeostasis and disease.

## Experimental Procedures

### Resource Availability

#### Lead contact

Further information and requests for resources and reagents should be directed to and will be fulfilled by the lead contact, Kaelyn Sumigray

#### Model system and permissions

All studies were conducted with Institutional Animal Care and Use Committee (IACUC) approval from Yale University’s Standing committee on animals, and in accordance with institutional guidelines for the humane treatment of animals.

#### Experimental animals

Male adult Sprague Dawley rats (*Rattus norvegicus*) with an average weight of 200 g were purchased from Charles River. Rats were housed in the animal care facility of the Yale Animal Resources Center according to institutional guidelines for the care and use of laboratory animals.

#### Cell dissociation and droplet-based sequencing

Epithelial cell suspensions were generated as described previously with modifications (Sumigray et al., 2018). Briefly, the proximal jejunum of adult male Sprague Dawley rats was isolated and rinsed in ice-cold PBS. The tissue was segmented into ∼2 cm fragments and opened longitudinally to expose the epithelium. The tissue was incubated in 30 mM EDTA in HBSS (Gibco, 14170112) at 37°C for 20 min. The tissue was vigorously shaken to release epithelium. The epithelium was collected into a 15 mL conical tube, incubated on ice, and allowed to settle via gravity. The epithelium was washed twice in HBSS, then incubated in 1 mg/ml dispase II (Roche, GE-0619) in HBSS for 10-12 minutes at 37°C with frequent shaking. A total of 1 mL FBS and 2 µL DNase I was added to the cell suspension, and the cells were passed through 70 µm strainers. Cells were collected by centrifugation at 300 G for 5 min at 4°C. Cell pellets were washed with HBSS + 10% FBS and centrifuged again at 300 G for 5 min at 4°C. Cells were resuspended in HBSS + 10% FBS at a 1200 cells/µL. Prior to droplet-based scRNA-seq, cells were run through a 40 µm filter. Single cell suspensions were processed through several additional rounds of purification, centrifugation and resuspension before the generation of an emulsion using the NextGem v3.1 3’UTR capture kit (10x Genomics) with a target recovery of 10,000 cells. The resulting reverse transcription and generation of the sequencing library were performed according to the manufacturer’s protocols. The final library was sequenced to a target recovery of 300 M read pairs and conducted at the Yale Center for Genomic Analysis.

#### Analysis of rat scRNA-seq data

##### Pre-processing

Sequencing data were demultiplexed and processed using CellRanger v5.0 with alignment to Rnor6, followed by clustering and expression analysis using Seurat v.4.0.1 package in R (v.4.0.3). Prior to downstream processing, we recovered 20,285 cells with a median transcript count of 32,331 UMI per cell. Cells with > 200 and < 5800 genes and > 5000 transcripts were retained. Cells with > 40% mitochondrial transcripts were excluded. Libraries from two separate rats were integrated. Data were normalized and log-transformed using the LogNormalize function in Seurat. All clusters expressing Cd45 (Ptprc) or hemoglobin (Hba-a1) were removed. This left a total of 7,362 cells, confirmed to be epithelial by Epcam expression. Clusters were assigned to known cell types by signature gene markers, as outlined in Supplemental Table 1.

#### Dimensionality reduction and clustering

The top 2000 variable genes were used for principal component analysis, and the top 20 principal components were selected and visualized by UMAP. We identified clusters using the “FindClusters” function in Seurat, using the parameter resolution 0.3. As mentioned above, clusters enriched for Ptprc or Hba-a1 were removed, and data were rescaled. The “FindNeighbors” and “FindClusters” functions were rerun, with 20 principal components and a resolution of 0.5.

The AddModule Score function in Seurat was used to generate scores for cell type markers and for “Notch-on” and “Notch-off” scores. The genes that contributed to each score are listed in Supplemental Table 1.

#### Rat Intestinal Organoid Derivation and Cell Culture

Rat intestinal organoids were established from the crypts of rat proximal jejunum as described previously (Zagoren *et al*., 2023). Briefly, following euthanasia, rat proximal jejunum was isolated and opened longitudinally to expose villi, which were scraped off with a glass microscope slide. Intestinal fragments were incubated 30 min in 3 mM EDTA at 4°C, after which the intestinal segment was shaken vigorously in PBS with fine tweezers under a dissecting microscope until the PBS primarily contained crypts. Crypts were pelleted at 250 *g* and the crypt pellet was added to growth factor-reduced Matrigel. Rat intestinal organoids were then cultured in rat intestinal organoid media (Zagoren *et al*., 2023) at 37°C, 5% CO_2_ and passaged via mechanical disruption every 3-5 days as needed.

#### Immunofluorescence staining

For cryosections, 10 µm-thick sections of rat proximal jejunum or distal duodenum were fixed in 4% paraformaldehyde (PFA) in PBS with 0.2% TritonX-100 (American Bio, AB02025) for 8 minutes. Tissues were blocked in 3% bovine serum albumin (Sigma, A9647) and 5% normal donkey serum (Sigma, D9663) for 1 hr. Primary antibody was then added for 15 min-overnight (O/N), depending on the antibody. After washing three times with PBST, sections were incubated in secondary antibody for 10 minutes at RT. Sections were then washed three additional times in PBST, after which they were mounted in Antifade and imaged on an upright Zeiss AxioImager with Apotome 2 attachment (Zeiss, Germany) with a Zeiss AxioCam 506 mono camera. Objectives used were Plan Apochromat 10x/0.45 air, 20x/0.8 air, 40x/1.3 oil, 63x/1.4 oil.

For whole mount staining, organoids were fixed in 4% PFA in PBST in the cell culture dish for 10 minutes. Organoids were released from the Matrigel by pipetting up and down, collected in a 1.5 mL tube, and PFA was removed and replaced with PBST. After allowing organoids to settle, the pellet was resuspended in block and incubated at room temperature for 45 minutes. Primary antibody incubation was at RT for 45min-O/N, depending on the antibody. Organoids were then washed three times in PBST, allowing them to settle at the bottom of the tube prior to each wash. Secondary antibody was added for 30 minutes at RT. Organoids were again washed three times in PBST. Lastly, organoids were mounted in Antifade and imaged on a Leica Stellaris 5 confocal laser scanning microscope with a white light laser using a HC FLUOTAR 25X/0.95 W VISIR objective or HC PL APO 40X/1.10 W CORR CS2 objective (Leica, Germany).

The primary antibodies used were as follows: Rabbit anti-CFTR antibody, used 1:500 O/N, was described previously (Golin-Bisello et al., 2005). Rabbit anti-Best4 antibody, used 1:250 O/N, is a custom rabbit polyclonal antibody synthesized against the Best4 extracellular domain (290-454 aa) and purified by Genscript for the purposes of this study. The remaining primary antibodies are commercially available: Mouse anti-Meis1 (1:100 O/N, Invitrogen, MA5-27191), Mouse anti-Prox1 (1:100 O/N, Novus Biologicals, NBP1-30045), Goat anti-Notch2 (1:100 15min, R&D Systems, AF1190-SP), Rabbit anti-Muc2 (1:1000 O/N, Abcam ab272692), Rabbit anti-Dclk1 (1:1000 O/N, Abcam ab31704), Rabbit anti-ChgA (requires prefixation, 1:500 O/N, Abcam, ab254557), Sheep anti-Dll1 (1:100 O/N, R&D Systems, AF5026-SP), Rabbit anti-Ki67 (1:200,1hr, Abcam, ab15580), Rabbit anti-Phospho Histone3 (1:500, 1hr, Cell Signaling, 3377), Rabbit anti-Hes1* (1:2500 Cell Signaling, 11988S) * requires 7.5 min incubation with Tyramide Signal Amplification Kit (ThermoFisher, B40922)

Secondary antibodies were used at 1:200 as follows, except for DAPI (5µg/mL, Invitrogen, D1306). Donkey anti-Goat AF488 (Jackson, 705-545-147), Donkey anti-Mouse AF488(Jackson, 715-545-151), Donkey anti-Rabbit AF488 (Jackson 711-545-152), Donkey anti-Goat AF647 (Jackson, 705-605-147), Donkey anti-Mouse AF647 (Jackson, 715-605-151), Donkey anti-Rabbit AF647 (Jackson, 711-605-152), phalloidin AF647 (Invitrogen, A22287), Donkey anti-Goat RRx (Jackson, 705-295-147), Donkey anti-Mouse RRx (Jackson, 715-295-151), Donkey anti-Rabbit RRx (Jackson, 711-295-152), Donkey anti-Sheep RRx (Jackson,713-295-147).

For colabeling of Best4 and CFTR, FlexAble Antibody labeling kits CoraLite Plus 488 and CoraLite Plus 555 were used according to the manufacturer’s protocols (Proteintech).

#### Animal treatment with DAPT

Adult male Sprague-Dawley rats (n=2 for DAPT treatment, n=2 for DMSO control) were injected with 20 mg/kg DAPT or an equivalent volume of DMSO 1x daily for five consecutive days. At day 6, rats were sacrificed in accordance with IACUC-approved institutional guidelines. The intestine was dissected following a previously described protocol (Zagoren *et al*., 2023) and distal duodenum and proximal jejunum were frozen in TissueTek OCT Compound (Sakura, 4583) for cryosectioning.

For organoids: DAPT powder (Millipore, 565770) was dissolved in DMSO to a stock concentration of 10 mM. Rat jejunum organoids were incubated in 5 µM DAPT or an equivalent volume of DMSO control for 3 days at 37°C, 5% CO_2_.

#### Forskolin Swelling of Organoids

Differentiated rat intestinal organoids (∼2-3 days post-passage) were cultured in rat intestinal organoid media in 35 mm glass-bottom dishes. For treated organoids, forskolin (Sigma, F3917), was diluted to a 10 µM working concentration, while for control organoids an equivalent volume of DMSO (vehicle) was added. Swelling was imaged every 10 minutes for 120 minutes total using the brightfield setting of a Leica Stellaris 5 confocal microscope.

## Statistical Analysis

All results are expressed as mean values ± standard deviation. The significance of the differences between different treatments and time points was compared using a two-tailed unpaired t-test in Prism 9 (GraphPad) unless noted otherwise in Figure legend. *p* values less than 0.05 (*p* < 0.05) were considered statistically significant. For all plots, error bars indicate SD.

## Acknowledgments

We thank members of the Sumigray and Ameen labs for thoughtful discussion and for comments on the manuscript, and the Smith lab for discussion of computational analysis. Research reported in this publication was supported by the National Institute Of Diabetes And Digestive And Kidney Diseases of the National Institutes of Health under Award Number F31DK137452 (EZ), the Yale Predoctoral Training Program in Genetics (5T32GM007499-43), as well as a Charles H. Hood Foundation Child Health Grant (KS), a Cystic Fibrosis Foundation grant (004741P222) to KS, and the National Institute of General Medical Sciences of the National Institutes of Health under Award Number 1R35GM150645-01 (KS).

## Author contributions

EZ, NAA and KS designed the study. EZ, NAA, ZDS, and KS performed experiments. EZ, ND and KS performed data analysis. EZ, ZDS, and KS wrote the manuscript.

---

**Figure S1. Cluster identification by marker gene expression**

A-H) UMAP of single cell RNA-seq data from rat intestinal jejunum color-coded by expression of A) stem cell markers, B) cell cycle score, C) enterocyte markers, D) Goblet cell markers, E) Paneth cell markers, F) EEC markers, G) Tuft cell markers, H) CHE markers. Marker genes are listed in Supp. Table 1. I) Dotplot for expression of differentiated cell type markers and enterocyte-associated genes across intestinal epithelial cell types. CHEs did not express markers of any other differentiated cell types or enterocyte-associated genes. J) Heatmap illustrating cell-type specific characteristic gene expression patterns.

**Figure S2. Secretory cell cluster characterization**

A-G) UMAP of reclustered stem and secretory progenitors color-coded by expression of A) stem cell markers, B) cell-cycle score, C) Goblet cell markers, D) Paneth cell markers, E) EEC markers, F) Tuft cell markers, G) CHE markers. Marker genes are listed in Supp. Table 1. H) UMAP of stem and secretory cell clusters color-coded by Dll1 expression. I) Stem and secretory cell UMAP annotated by cell cycle stage: S-phase (red), G1 (green), G2/M (blue). J) Secretory progenitor in rat proximal jejunum, occupying the +4 position and expressing high levels of membrane-localized Dll1 (magenta). DAPI in white. Scale bar, 10 µm. K) Dll1 (magenta) and Ki67 (cyan) costaining of +4 cells. Scale bar, 10 µm. L) Dotplot from scRNA-seq data of expression of transcription factors enriched in CHEs. M) Best4+ colon cells (Best4, cyan) were Prox1+ (magenta) in rat proximal colon. DAPI in white. N) Meis1 (magenta) was undetectable in Best4+ colon cells (Best4, cyan). DAPI in white. Scale bar for M-N, 20 µm.

**Figure S3. Rat intestinal organoids generate CHEs**

A) Undifferentiated rat jejunum organoid form spheroids. Scale bar, 50 µm. B) CFTR (yellow) in undifferentiated rat intestinal organoids. Phalloidin in magenta. Scale bar, 20 µm. C) Differentiated rat intestinal organoid ∼3 days post-passaging. Scale bar in panel A. D) Rat organoids differentiate to form CHE cells. CFTR (yellow), phalloidin (magenta). Scale bar, 20 µm. E) Distribution of CFTR in organoids recapitulates *in vivo* CFTR distribution. F) CFTR intensity in organoids recapitulates *in vivo* distribution. two-tailed paired t test. *p*<0.0001. G-J) CHE in organoids also expressed Meis1, Prox1, Best4, and Notch2 as seen *in vivo*. G) CHEs with CFTR (cyan) and Meis1 (magenta). H) Best4+ cells (yellow) in organoids. Phalloidin in magenta. I) CHEs that are Prox1+ (white) and Notch2+ (magenta). J) CHEs that are CFTR+ (yellow) and Notch2+ (cyan). Scale bars, 20 μm.

**Table S1.** Genes used to generate module scores for cell cluster identification and Notch signaling scores.

| **Module Score** | **Gene** |
| --- | --- |
| Stem cell | Lgr5, Ascl2, Slc12a2, Axin2, Olfm4, Gkn3 |
| Cell cycle | Mki67, Cdk4, Mcm5, Mcm6, Pcna |
| Enterocyte | Alpi, Apoa1, Apoa4, Fabp1 |
| Goblet | AABR07006030.1, Tff3, Agr2 |
| Paneth | Lyz2, Wnt3, Dll4, Defa24, Wnt11 |
| Enteroendocrine | Chga, Chgb, Tac1, Tph1, Neurog3 |
| Tuft | Dclk1, Trmp5, Gfi1b, Il25 |
| CHE | Cftr, Best4 |
| Notch on | Notch1, Notch2, Notch3, Notch4, Hes1, Nrarp |
| Notch off | Dll1, Dll4, Jag1, Jag2, Atoh1 |

