## Supplementary figures and images for "A second wave of Notch signaling diversifies the intestinal secretory lineage"

### Supplemental Figures 1-3

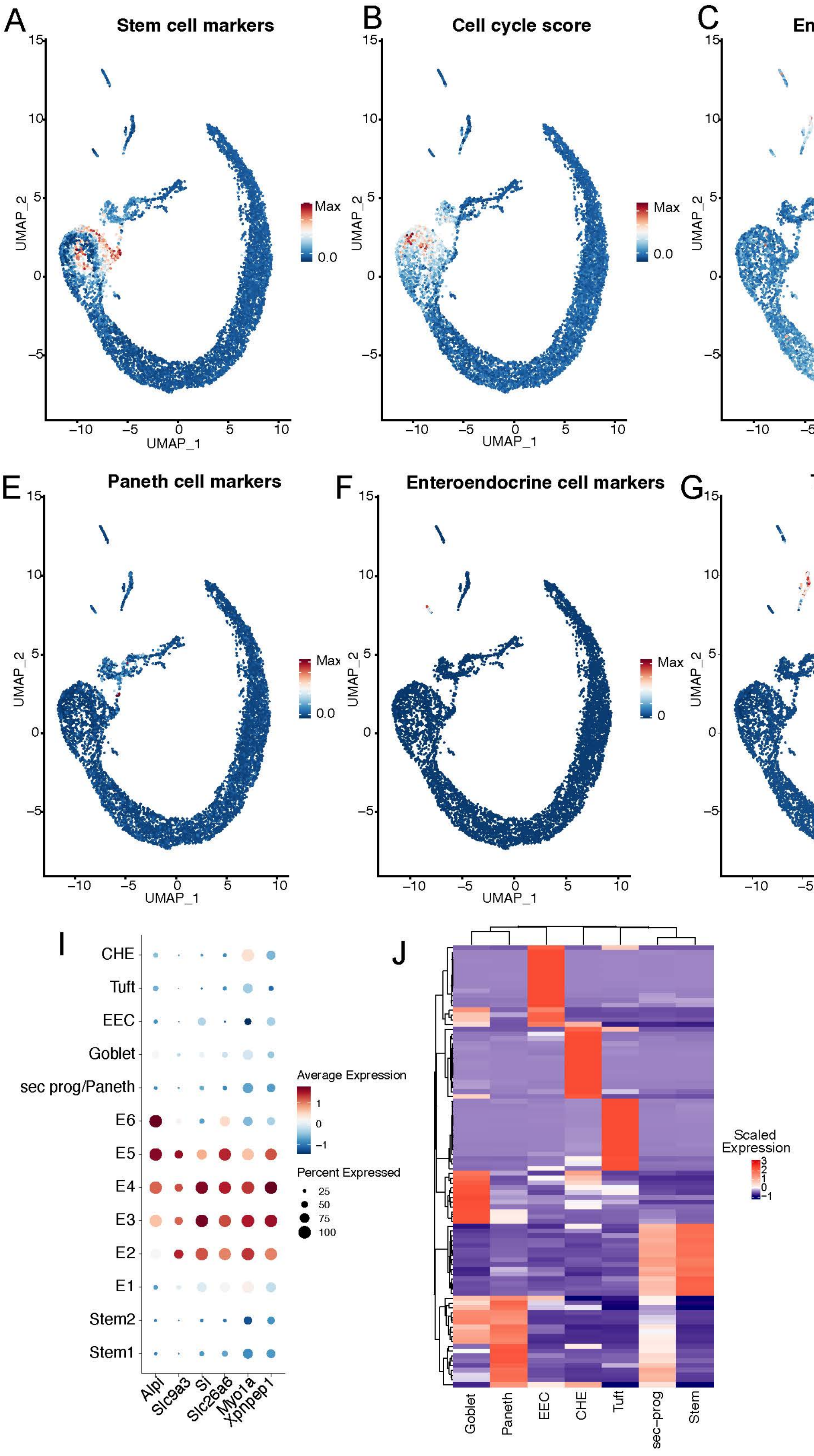

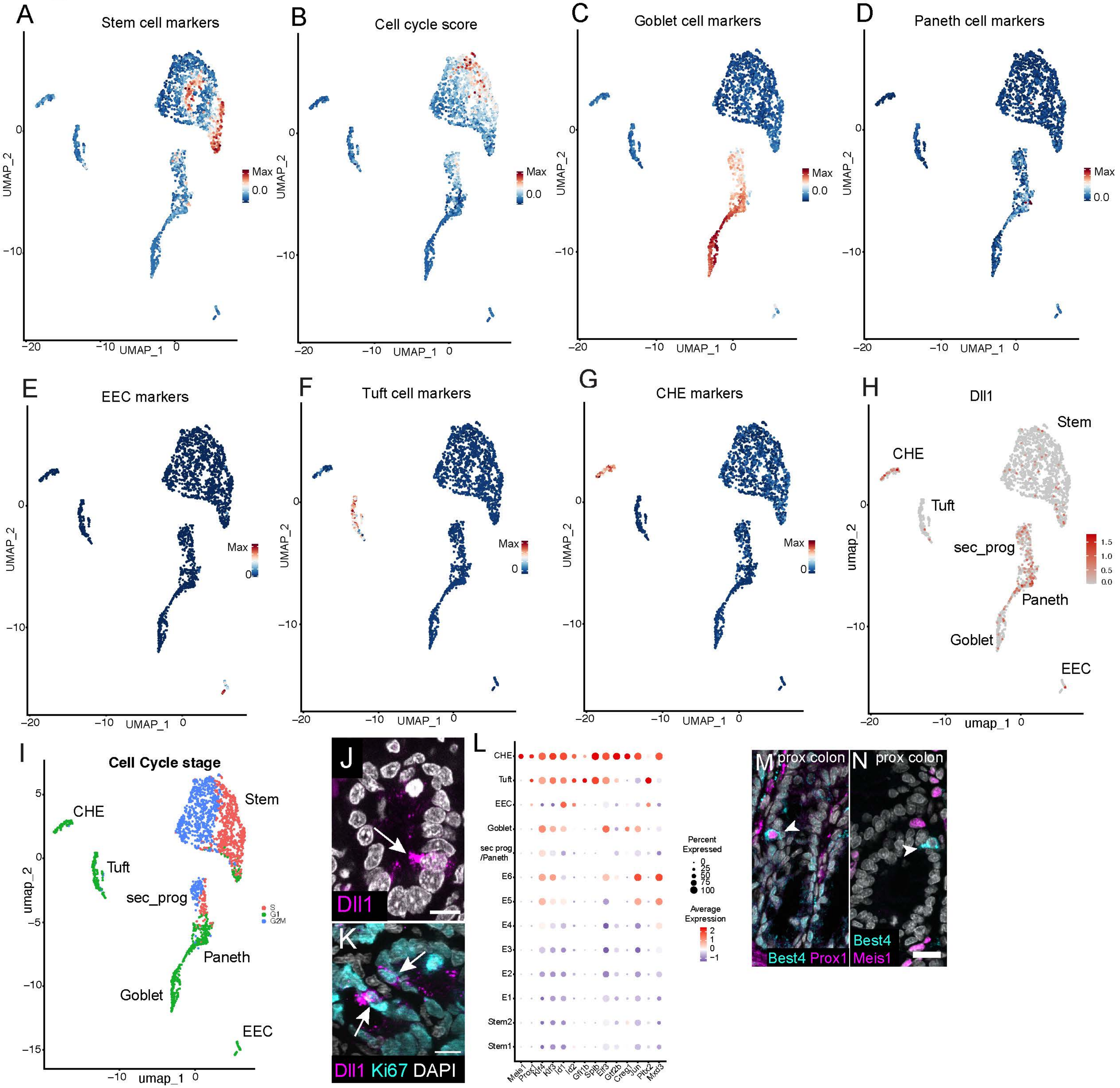

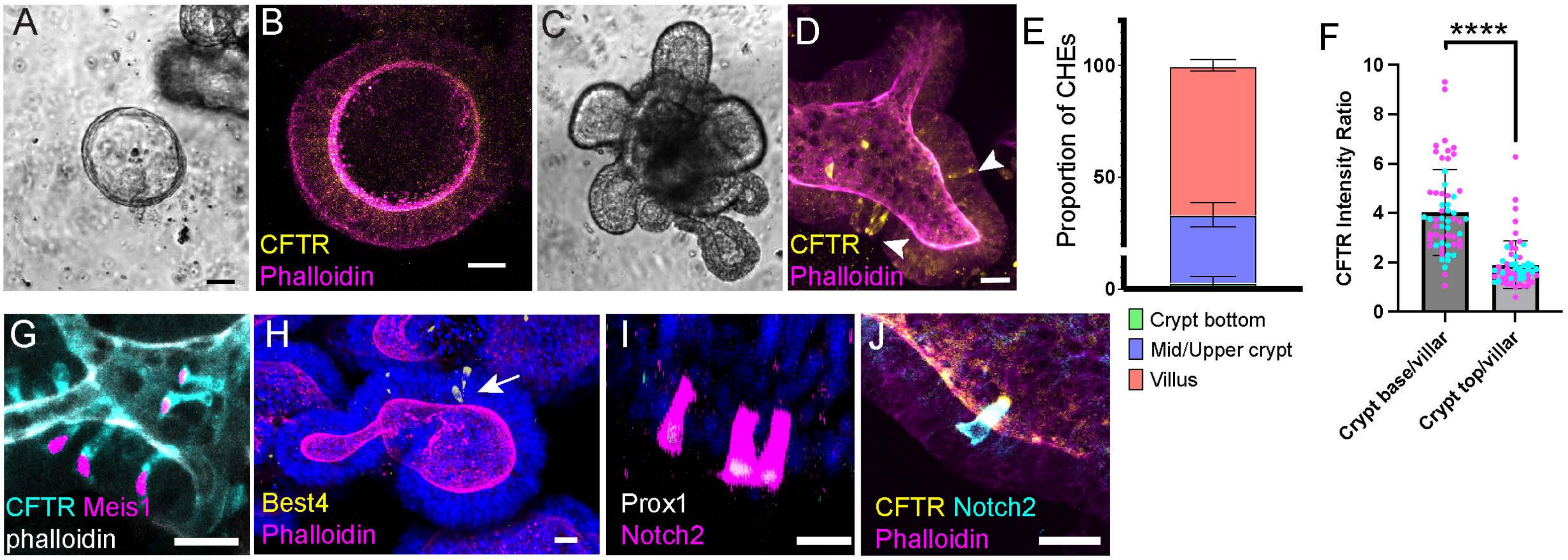
